# Metabolic alterations in a *Drosophila* model of Parkinson’s disease based on *DJ-1* deficiency

**DOI:** 10.1101/2021.12.15.472768

**Authors:** Cristina Solana-Manrique, Francisco José Sanz, Isabel Torregrosa, Martina Palomino-Schätzlein, Carolina Hernández-Oliver, Antonio Pineda, Nuria Paricio

## Abstract

Parkinson’s disease (PD) is the second most common neurodegenerative disorder whose physiopathology is still unclear. Besides, it is urgent to discover new biomarkers and therapeutic targets to facilitate its diagnosis and treatment. Previous studies performed in PD models and samples from PD patients already demonstrated that metabolic alterations were associated with this disease. In this context, the aim of this study is to give a better understanding of metabolic disturbances underlying PD pathogenesis. To achieve this goal, we used a *Drosophila* PD model based on inactivation of the *DJ-1β* gene (ortholog of human *DJ-1*). Metabolomic analyses were performed in 1-day-old and 15-day-old *DJ-1β* mutants and control flies using ^1^H nuclear magnetic resonance spectroscopy, combined with expression and enzymatic activity assays of proteins implicated in altered pathways. Our results showed that PD model flies exhibit protein metabolism alterations, a shift from tricarboxylic acid cycle to glycolytic pathway to obtain ATP, together with an increase in the expression of some urea cycle enzymes. Thus, these metabolic changes could be contributing to PD pathogenesis and might constitute possible therapeutic targets and/or biomarkers for this disease.

## INTRODUCTION

In the last 25 years, neurodegenerative diseases (NDs) have become an important challenge for general health, especially in older population groups, and are expected to grow in the future due to an increase in life expectancy [1]. Among them, Parkinson’s disease (PD) is the most common motor disorder, affecting more than 1% of the population over 60 years. PD is caused by the selective and progressive loss of dopaminergic (DA) neurons in the *substantia nigra pars compacta* (SNpc), which results in striatal dopamine deficiency and leads to disturbances in motor, autonomic and psychiatric functions [2–4]. DA neuron loss is accompanied, in some cases, by the presence of α-synuclein aggregates in the surviving neurons, known as Lewy bodies [2,4]. The exact mechanism behind DA neurodegeneration in PD is still unclear, being probably a combination of genetic predisposition and environmental factors [2,5]. Indeed, multiple pathways and mechanisms seem to participate in PD pathogenesis like accumulation of misfolded protein aggregates, mitochondrial dysfunction, increased oxidative stress (OS), energy failure, neuroinflammation, and genetic mutations [6]. Interestingly, recent studies have shown that bioenergetic alterations may play a key role in PD neuropathology. Concretely, an increase in the glycolytic rate has been observed in PD models, suggesting that there is a link between glucose metabolism, cellular bioenergetics, redox homeostasis and neuronal death [7,8].

Most PD cases are idiopathic (iPD), whose aetiology is multifactorial. However, 5-10% of PD patients suffer from monogenic forms of the disease caused by highly penetrant mutations that are family-linked [2,5]. In the last years, mutations in several genes have been associated with these familial forms of PD (fPD), allowing to discover different mechanisms underlying PD pathogenesis [5,9]. Indeed, some of the proteins encoded by these genes are involved in a set of molecular pathways that upon perturbation trigger a neuropathology that resembles to, or is clinically indistinguishable from iPD, except for the age at onset [10]. Among them, *DJ-1* is a causative gene for fPD [11] that was initially described as an oncogene. Nevertheless, other functions have been ascribed to the DJ-1 protein such as transcriptional regulation, chaperone and protease activity, scavenger of reactive oxygen species (ROS) or mitochondrial homeostasis [12–14]. Remarkably, an over-oxidized and inactive form of the DJ-1 protein has also been found in brains of iPD individuals suggesting that it may play a central role in the development of the disease [15]. Therefore, results obtained in PD models with impaired DJ-1 function could be also applicable to human iPD forms.

Currently, PD diagnosis is limited and based on the detection of classic motor symptoms that appear when about 60-80% of SNpc DA neurons are lost, after many years of ongoing disease [16]. Thus, it is urgent to find simple, useful and low-cost biomarkers that allow an early PD diagnosis to address its progression in the initial stages of the disease [17]. Biomarkers are also needed for distinguishing different PD types, predicting the course of the disease, or monitoring the effect of disease-modifying therapies [18,19]. Metabolomic technologies allow the analysis of a great amount of low-molecular-weight molecules, offering an overview of the molecular complexity of a biological system and the metabolic pathways that can be altered in a pathological state [20,21]. Therefore, metabolomic studies are being carried out to identify new biomarkers in PD and other NDs [22,23] as well as therapeutic targets and metabolic alterations implicated in PD pathogenesis [23].

*Drosophila melanogaster* has emerged as an important model organism in the study of PD physiopathology. In this scenario, several genetic-based and chemically-induced PD models have been developed in *Drosophila* [24,25]. Among them, flies harboring mutations in *DJ-1β* (ortholog of *DJ-1* human gene) present typical PD phenotypes, like motor impairment, and increased levels of OS markers [26–28]. In addition, *DJ-1β* mutants exhibit increased activity of key glycolytic enzymes [7]. Given the great complexity and heterogeneity in the metabolome of human samples as well as their limited availability and obtaining, simple models such as *Drosophila* to study PD-associated metabolome-wide alterations are becoming very useful [29–31]. In fact, metabolomic studies have been performed in different *Drosophila* models of NDs. For instance, a metabolomic analysis in a *Drosophila* model of Charcot-Marie-Tooth disease based on *GDAP1* deficiency allowed to identify the involvement of insulin signaling in the characteristic neuromuscular degeneration of this disease [32]. Regarding PD, Shukla *et al* [21] performed a metabolomic analysis in a paraquat-induced *Drosophila* model of iPD, in which altered levels of amino acids, lipids and carbohydrates were identified [21]. In such a scenario, we aimed to find new pathways underlying PD pathogenesis and/or new potential biomarkers by carrying out a metabolomic analysis in *DJ-1ß* mutant flies at different ages. Changes in metabolite levels in *DJ-1β* mutants allowed us to identify metabolic alterations in PD model flies, such as changes in amino acids metabolism, a switch from tricarboxylic acid (TCA) cycle to glycolysis and other pathways affected like the urea cycle (UC). To our knowledge, this is the first metabolomic study carried out so far in a *Drosophila* fPD model.

## MATERIALS AND METHODS

### *Drosophila* strains

All stocks and crosses were cultured on standard *Drosophila* food at 25°C. For this work, *DJ-1β*^*ex54*^ flies (hereafter called *DJ-1β* mutant) were used as PD model flies [33] and *y*^*1*^,*w*^*1118*^ were used as controls.

### Metabolite extraction

Metabolite extraction was performed as previously described in [32]. Twelve samples of 1-day-old and 15-day-old individuals were prepared for each batch of *DJ-1β* mutant and controls, containing fifteen female flies each. They were frozen in a microtube by immersion in liquid nitrogen. Then, 240 µl of ice-cold methanol and 48 µl of ice-cold Milli-Q water were added to each vial. After 5 min, flies were homogenized with a small mortar for 60 s. Samples were vortexed and 120 µl of ice-cold CHCl_3_ and 120 µl of ice-cold Milli-Q water added and vortexed again. After 15 min at 4°C, samples were centrifuged at 10000*g* during 15 min at 4°C. The resulting two phases (upper phase: polar metabolites, lower phase: non-polar metabolites) were separated. Solvents from the polar phase were eliminated by freeze-thawing and samples were stored at −80°C until measurement.

### NMR analysis

Metabolite extracts were allowed to thaw during 5 min at 4°C and dissolved in 550 µl of NMR buffer [0.1 M phosphate buffer pH 7.4 in D2O, with 0.1 mM 3-(Trimethylsilyl)propionic-2,2,3,3-d4 acid sodium salt (TSP) as internal standard] for the polar phase. Polar samples were analyzed on a 600 MHz Bruker NMR spectrometer equipped with a cryoprobe using a 1D noesy experiment including presaturation for water signal suppression and a 50 ms mixing delay. Spectra were acquired with 512 scans, a relaxation delay of 4 s and a spectral width of 18028 Hz and processed with an exponential line broadening factor of 0.5. For metabolite identification, 2D TOCSY and HSQC experiments were acquired for selected samples. All experiments were acquired at 37°C.

### Metabolite assignment and quantification

NMR spectra were processed and analyzed with NmrProcflow [34]. Baseline correction was performed with local level 5 and spectra alignment with the option “least square”. Bucketing was applied manually with a SNR of 3, resulting in 310 buckets. The signals corresponding to methanol, chloroform, TSP and H_2_O were excluded. Data were exported from NMRProcFlow tool and integration values were normalized to total intensity (CSN) to minimize variability. For the statistical analysis, a multivariate data analysis was performed using SIMCAP 12.0 Software (Umetrics, Sweden) with the normalized integral values. First, spectra were normalized with Probabilistic Quotient Normalization. Principal component analysis (PCA) was performed for a first overview to evaluate clustering trends between samples and to identify outliers. Secondly, Orthogonal Partial Least Squares Discriminant Analysis (OPLS-DA) was performed for discrimination analysis between groups. Furthermore, the analysis of S-plots allowed to define metabolites that were essential form the discrimination between groups. OPLS-DA models were validated by permutation. In a second step, a pathway analysis was applied using the web-based software for metabolomics, MetaboAnalyst 5.0 Software (https://metaboanalyst.ca/).

### Measurement of ATP levels

ATP levels were measured using the ATP Determination Kit (Invitrogen) following manufacturer’s instructions. Briefly, groups of five 15-day-old *DJ-1β* and control female flies were homogenized in 200 µl of reaction buffer (supplied by the commercial kit). Then, fly extracts were boiled 4 min and centrifuged at 18.500*g* for 10 min at 4°C in order to discard debris. Subsequently, 5 µl of fly extracts were added to 100 µl of the standard reaction solution in a white 96-well plate and luminescence was measured using an Infinite 200 PRO reader (Tecan). All experiments were performed in triplicate and results are expressed as relative luminescence intensity per mg of protein normalized to control flies.

### Enzymatic activity assays

Protein extracts were obtained from groups of 20 *DJ-1β* mutants and control female flies as described in [7]. Aconitase (Aco; EC 4.2.1.3) activity was measured with the Aconitase Activity Assay Kit (#MAK051; Sigma-Aldrich), following manufacturer’s instructions and assays were performed in triplicate. Succinate dehydrogenase (SDH; EC 1.3.5.1) activity was measured using a protocol adapted from [35]. First, mitochondrial extracts were obtained from groups of 30 female *DJ-1β* mutants and control flies as described in [36]. Then, 50 µl of mitochondrial extracts were added to 50 µl of assay buffer [4 mM sodium azide, 50 µM 2,6-dichlorophenolindophenol and 2 µg/ml rotenone] and transferred to a 96-well plate. The reaction was started by adding succinate to a final concentration of 10 mM. Absorbance was measured at 600 nm using an Infinite 200 PRO reader (Tecan) every 30 s for 20 min at 25°C. Sample absorbance levels were measured, subtracting their corresponding blanks. Assays were performed in triplicate.

### RT-qPCR analyses

Total RNA from groups of ten 15-day-old *DJ-1β* mutants or control female flies was extracted and reverse transcribed as described in [7]. RT-qPCR reactions were performed as [7] and the following pairs of primers were used: *tubulin* direct primer (5’- GATTACCGCCTCTCTGCGAT −3’); *tubulin* reverse primer (5’- ACCAGAGGGAAGTGAATACGTG −3’); *arg* direct primer (5’- AGCTTTGACATCGACGCCTT-3’); *arg* reverse primer (5’- CTCCACGATGCTGATTCCCT-3’); *argL* direct primer (5’- CGCATCACATTATCGGTCGC-3’); *argL* reverse primer (5’- TCTCCCAGCGGCAAATCAG-3’).

### Statistical analysis and data representation

Data are expressed as means ± standard deviation (s.d.). The significance of differences between means was assessed using t-test except for NMR analysis where a different statistical analysis was performed (see section NMR analysis). Differences were considered significant when \**P* < 0.05. Data representations were performed using GraphPad Prims 6.0 Software (GraphPad Software, Inc., Chicago, IL, USA).

## RESULTS

### Impact of *DJ-1β* loss on the general metabolic profile

Previous studies showed that lack of *DJ-1* function leads to several metabolic alterations [7,37–39]. Indeed, we have recently demonstrated that *DJ-1β* mutant flies and *DJ-1*-deficient human neuroblastoma cells show an increase of the glycolytic pathway [7]. In order to identify additional metabolic changes caused by loss of *DJ-1β* function that could be contributing to PD pathophysiology, we undertook metabolomic analyses in *DJ-1β* mutants and control flies by NMR spectroscopy. They were performed in 1-day-old and 15-day-old flies, to detect early alterations that may constitute promising biomarkers for PD diagnosis as well as later ones that could help to evaluate disease progression.

Firstly, spectral peaks were obtained in all four experimental groups followed by the metabolite assignment of peaks that were statistically relevant. After this, the relative abundance of each metabolite in *DJ-1β* mutants and control flies at both ages was established to evaluate the differences between these groups. A multivariable statistical analysis was performed to find the most significant changes found in the metabolomic analysis, which are shown in Tables S1-S3. Metabolic alterations were characterized PCA to identify outliers and to evaluate the existence of any trend or aggrupation due to another variable. As shown in Figure 1 all samples grouped according to the phenotype and the specific age, especially *DJ-1β* mutant flies for which both ages are distinctly separated. This evaluation was followed by an OPLS-DA to establish the models that allow differentiating metabolic changes of all four experimental groups. In all comparisons, optimal models were obtained with R^2^ values close to 1 and reasonable predictive values (Q^2^) (data not shown).

**Figure 1.**
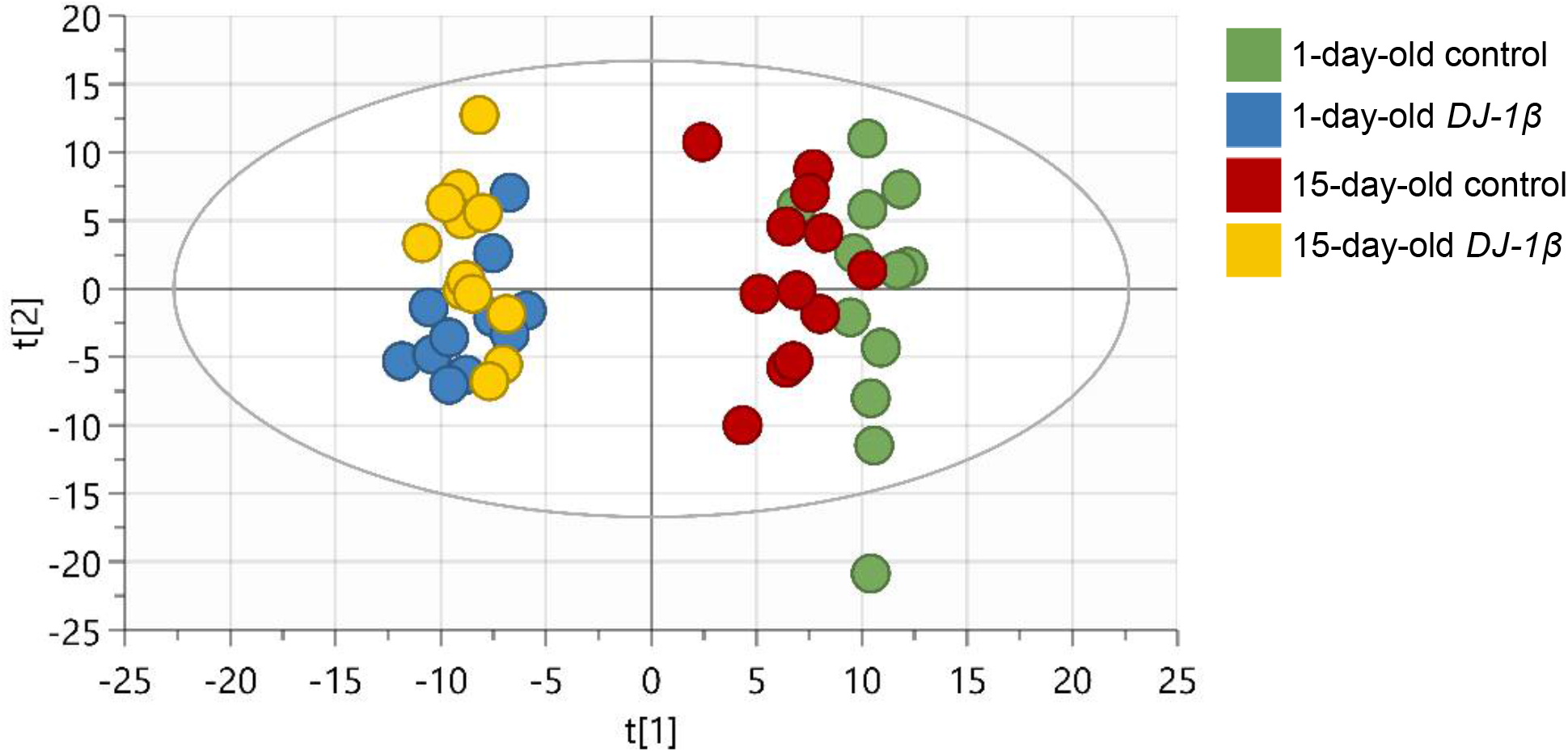
Metabolic profile of *DJ-1β* mutant flies compared to control. PCA-X score plot comparing all four experimental models: 1-day-old control flies (blue), 1-day-old *DJ-1β* mutant flies (green), 15-day-old control flies (yellow) and 15-day-old *DJ-1β* mutant flies (red).

To identify relevant pathways that could be perturbed in *DJ-1β* mutants at different ages, we used the MetaboAnalyst 5.0 software to examine metabolites showing significant alterations in such flies compared to controls (Tables S4-S6). As shown in Figures 2A-B and Tables 1-2, the most relevant changes with a considerable impact power between *DJ-1β* mutant and control flies at both ages belonged to several amino acid metabolism pathways, the TCA cycle (impact=0.24), pyruvate metabolism (impact=0.28) and glycolysis/gluconeogenesis pathways (impact=0.13). In addition, comparisons between 1-day-old and 15-day-old *DJ-1ß* mutant flies showed possible alterations in alanine, aspartate, and glutamine metabolism (impact=0.19), phenylalanine, tyrosine, and tryptophan biosynthesis (impact=0.50), phenylalanine metabolism (impact=0.38) and TCA cycle pathway (impact=0.24) (Figure 2C, Table 3). In summary, the most significant metabolic changes found in *DJ-1ß* mutants compared to control flies point to alterations in amino acid levels and their corresponding metabolism as well as in carbohydrate metabolism. Thus, components of these pathways may constitute therapeutic targets and/or potential biomarkers in PD. These findings constitute an initial step to further investigate genes, enzymes, or metabolites implicated in these pathways that could be contributing to PD physiopathology.

**Table 1.**
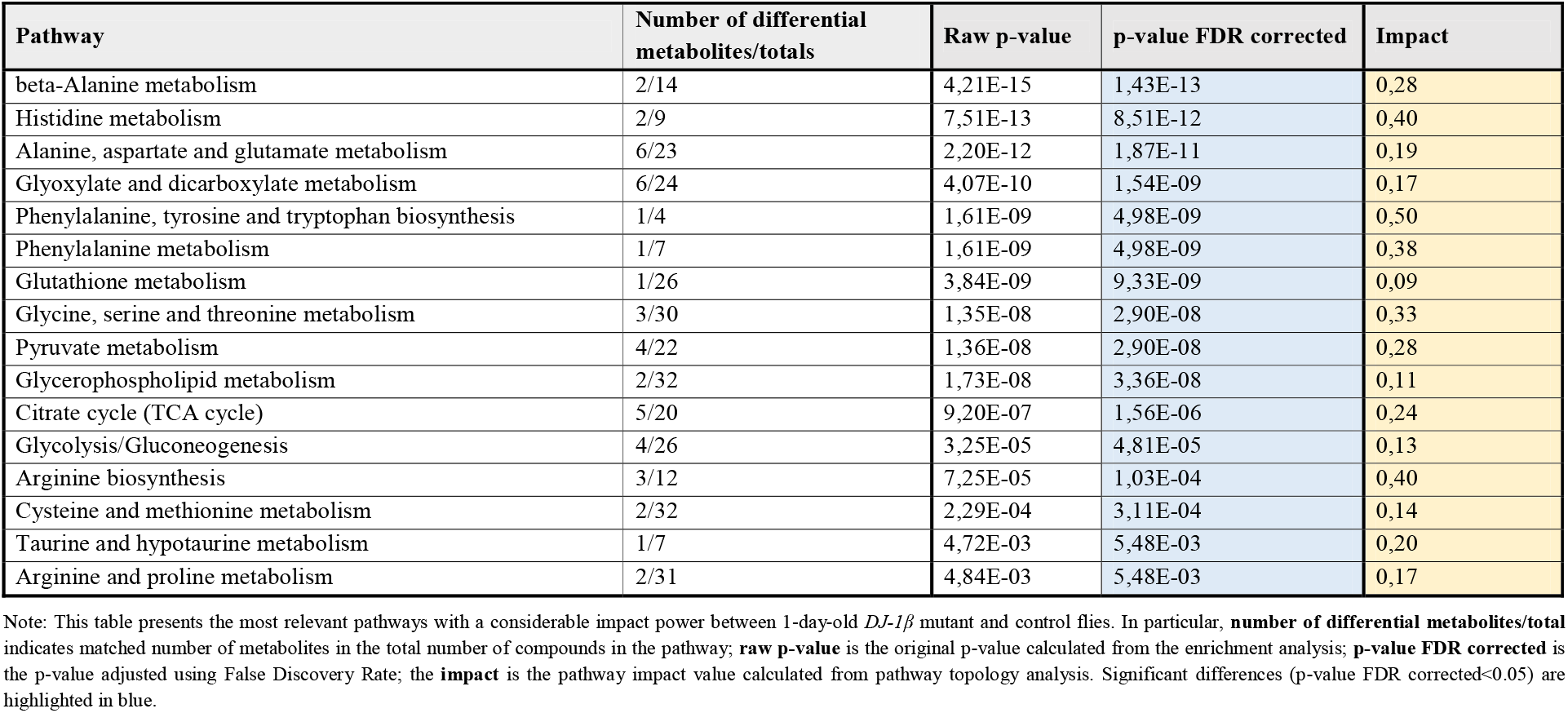
Selected results from the pathway enrichment analyses in 1-day-old *DJ-1β* mutants and control flies.

| Pathway | Number of differential metabolites/totals | Raw p-value | p-value FDR corrected | Impact |
| --- | --- | --- | --- | --- |
| beta-Alanine metabolism | 2/14 | 4,21E-15 | 1,43E-13 | 0,28 |
| Histidine metabolism | 2/9 | 7,51E-13 | 8,51E-12 | 0,40 |
| Alanine, aspartate and glutamate metabolism | 6/23 | 2,20E-12 | 1,87E-11 | 0,19 |
| Glyoxylate and dicarboxylate metabolism | 6/24 | 4,07E-10 | 1,54E-09 | 0,17 |
| Phenylalanine, tyrosine and tryptophan biosynthesis | 1/4 | 1,61E-09 | 4,98E-09 | 0,50 |
| Phenylalanine metabolism | 1/7 | 1,61E-09 | 4,98E-09 | 0,38 |
| Glutathione metabolism | 1/26 | 3,84E-09 | 9,33E-09 | 0,09 |
| Glycine, serine and threonine metabolism | 3/30 | 1,35E-08 | 2,90E-08 | 0,33 |
| Pyruvate metabolism | 4/22 | 1,36E-08 | 2,90E-08 | 0,28 |
| Glycerophospholipid metabolism | 2/32 | 1,73E-08 | 3,36E-08 | 0,11 |
| Citrate cycle (TCA cycle) | 5/20 | 9,20E-07 | 1,56E-06 | 0,24 |
| Glycolysis/Gluconeogenesis | 4/26 | 3,25E-05 | 4,81E-05 | 0,13 |
| Arginine biosynthesis | 3/12 | 7,25E-05 | 1,03E-04 | 0,40 |
| Cysteine and methionine metabolism | 2/32 | 2,29E-04 | 3,11E-04 | 0,14 |
| Taurine and hypotaurine metabolism | 1/7 | 4,72E-03 | 5,48E-03 | 0,20 |
| Arginine and proline metabolism | 2/31 | 4,84E-03 | 5,48E-03 | 0,17 |

Note: This table presents the most relevant pathways with a considerable impact power between 1-day-old *DJ-1 $\beta$* mutant and control flies. In particular, **number of differential metabolites/total** indicates matched number of metabolites in the total number of compounds in the pathway; **raw p-value** is the original p-value calculated from the enrichment analysis; **p-value FDR corrected** is the p-value adjusted using False Discovery Rate; the **impact** is the pathway impact value calculated from pathway topology analysis. Significant differences (p-value FDR corrected<0.05) are highlighted in blue.

**Table 2.**
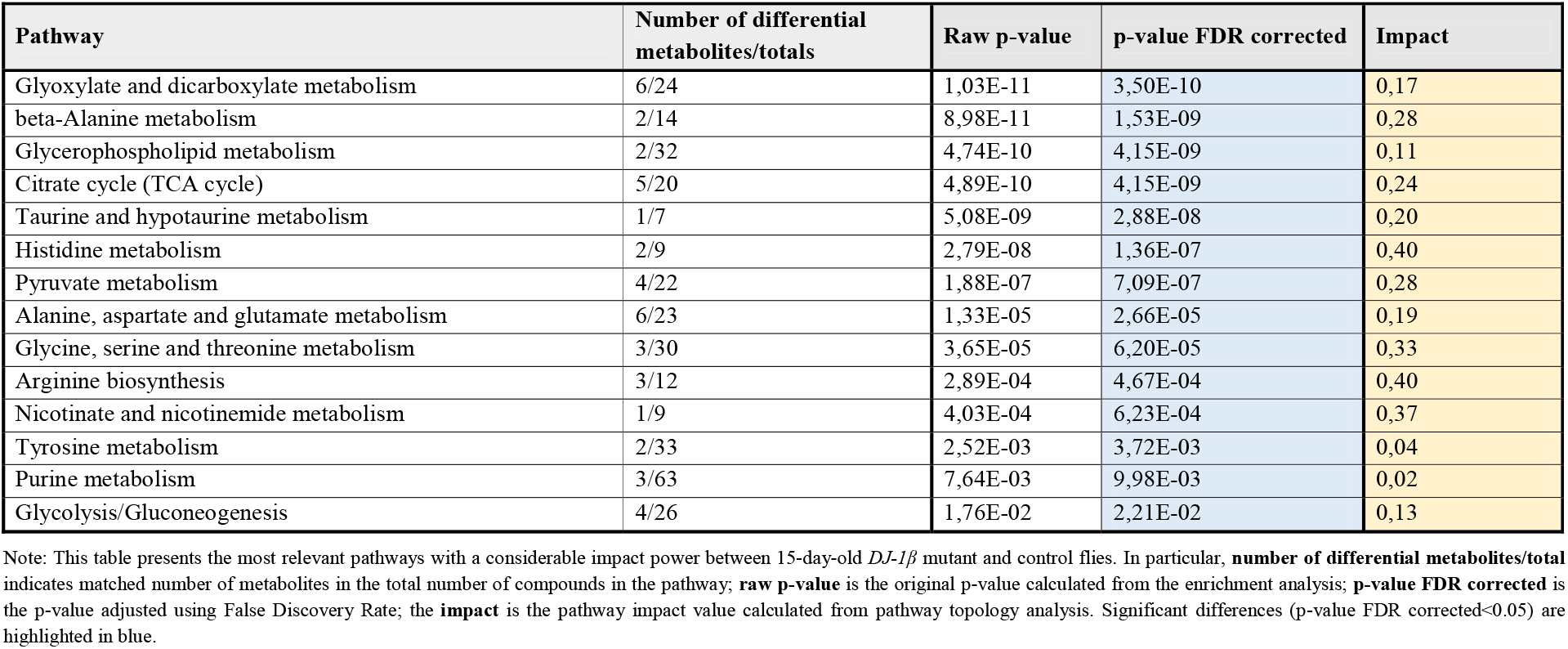
Selected results from the pathway enrichment analyses in 15-day-old *DJ-1β* mutants and control flies.

**Table 3.**
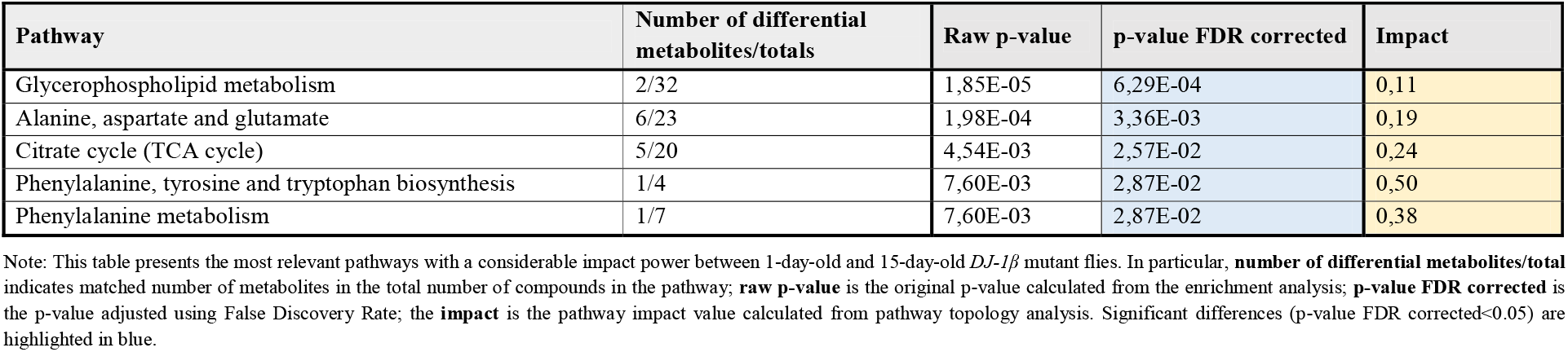
Selected results from the pathway enrichment analyses in 1-day-old and 15-day-old *DJ-1β* mutant flies.

**Figure 2.**
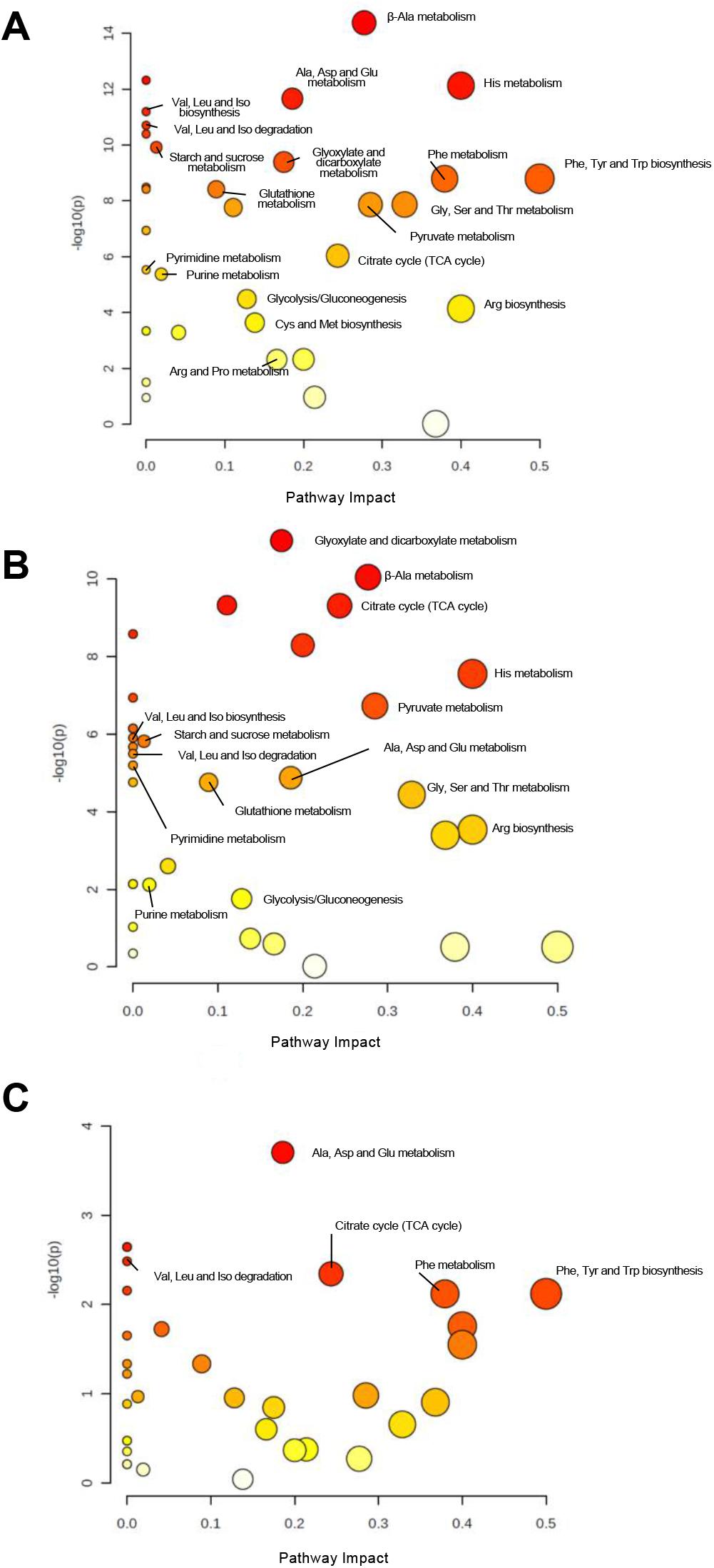
Pathway analysis reveals metabolic alterations in *DJ-1β* mutant flies. (A) Pathway enrichment analysis of differentially expressed metabolites between 1-day-old *DJ-1β* mutants and control flies. (B) Pathway enrichment analysis of differentially expressed metabolites between 15-day-old *DJ-1β* mutant and control flies. (C) Pathway enrichment analysis of differentially expressed metabolites between 1-day-old 15-day-old *DJ-1β* mutant flies.

### Alterations in amino acid content in *DJ-1β* mutant flies

In Figure 3A, a detailed overview of the differences in amino acids found between 1-day-old *DJ-1β* mutants and control flies is shown. There is a consistent reduction in the amounts of all the amino acids detected in the metabolomic analysis but β-alanine, acetyl-aspartate, N-acetyl-aspartate, asparagine, and glutamine, which are increased in 1-day-old *DJ-1β* mutants when compared to control flies. Similar results are found when comparing the amino acid profile of 15-day-old *DJ-1β* mutants and control flies with exception of glutamine, which showed no significant differences between both genotypes (Figure 3B). Among the amino acids whose levels are reduced in PD model flies we found the three branched-chain amino acids (BCAAs) (leucine, isoleucine and valine), which promote protein synthesis in the muscle [40], and essential amino acids (histidine, isoleucine, leucine, lysine, methionine, phenylalanine, threonine, tryptophan and valine). Interestingly, a reduction of both BCAAs and essential amino acids has been associated with an increase in the clinical severity of PD patients [41]. It is also noteworthy that there is a decrease in tryptophan levels in 1-day-old and 15-day-old *DJ-1β* mutant flies when compared to controls of the same age (Figures 3A-B). Tryptophan is an essential amino acid that participates in the kynurenine pathway to produce NAD^+^ and whose metabolism is altered in PD [42]. A further evaluation of the amino acid content led us to observe an increase in alanine, asparagine, glycine, isoleucine, leucine, phenylalanine and valine levels and a decrease in acetyl-aspartate in 15-day-old compared to 1-day-old *DJ-1β* mutant flies (Figure 3C). These results are consistent with those found in the amino acid profile in serum from patients with early or late PD, where alterations in alanine, arginine, phenylalanine and threonine were detected [43]. Therefore, these amino acids could be relevant in early PD diagnosis.

**Figure 3.**
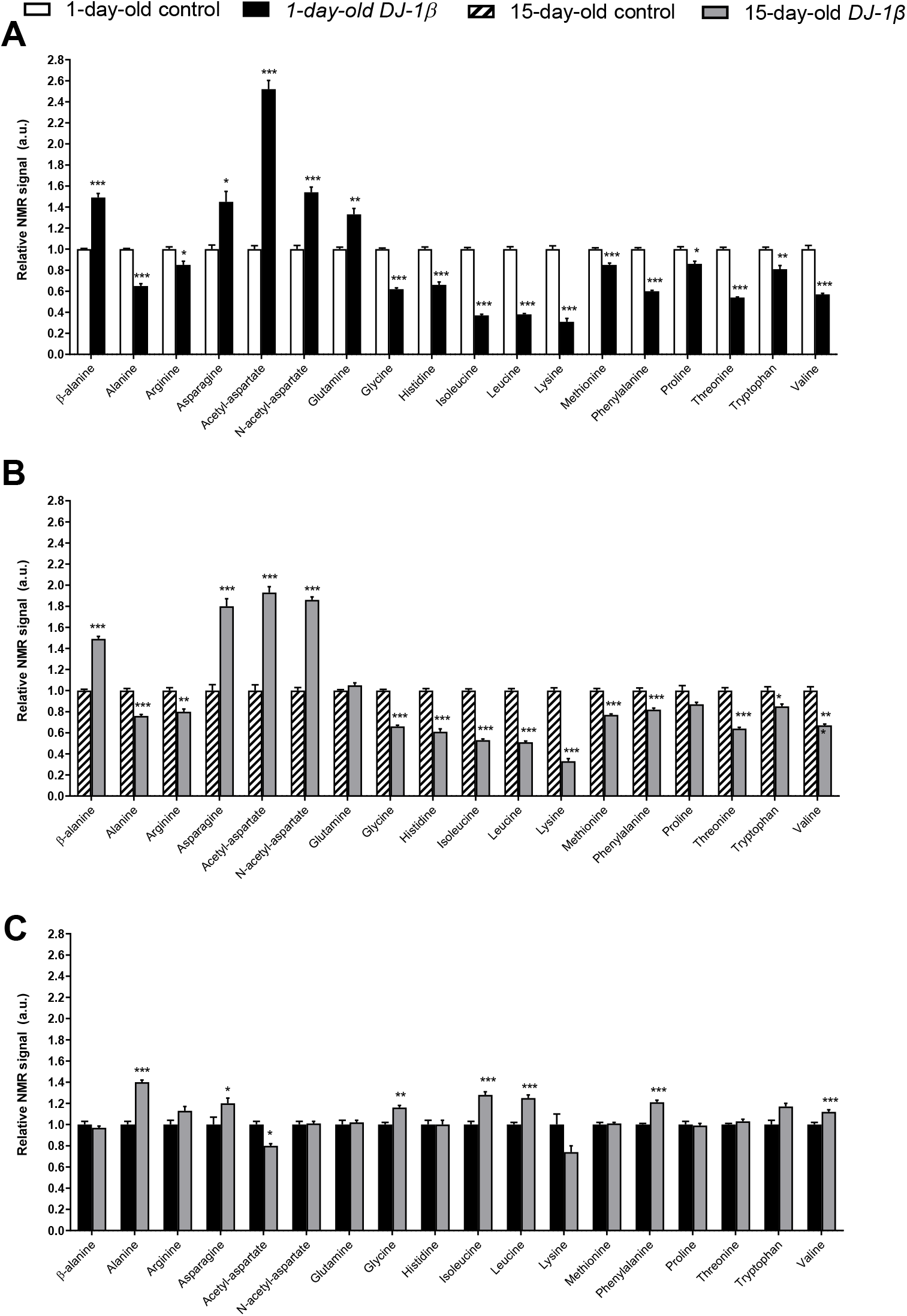
Alterations in amino acid content in PD model flies. (A) Relative NMR signals of amino acids comparing 1-day-old *DJ-1β* mutants and control flies. (B) Relative NMR signals of amino acids comparing 15-day-old *DJ-1β* mutants and control flies. (C) Relative NMR signals of amino acids comparing 1-day-old and 15-day-old *DJ-1β* mutant flies.

### *DJ-1β* deficiency leads to changes in carbohydrate metabolism

As mentioned above, *DJ-1β* mutant flies show alterations in the TCA cycle, pyruvate metabolism and glycolytic/gluconeogenesis pathways (Figure 2). These results support previous studies in which we demonstrated that PD model flies present an enhancement of glycolysis [7]. Therefore, we decided to delve into the study of carbohydrates metabolism, since several investigations have emphasized the importance of glucose usage in PD [44,45].

Regarding soluble sugars detected in the metabolic profiles, we found that 1-day-old *DJ-1β* mutants exhibit a decrease in glucose and pyruvate levels, and an increase in fructose and trehalose when compared to control flies of the same age (Figure 4A). An increase in trehalose is also observed in 15-day-old PD model flies, but changes in other sugars are not detected at that age, despite that a decrease trend in glucose and pyruvate levels is observed (Figure 4B). When comparing soluble sugar levels between 1-day-old and 15-day-old *DJ-1β* mutant flies, a reduction in fructose and an increase in glucose levels are observed in 15-day-old flies (Figure 4C), which could imply that glucose metabolism is reduced at late stages of PD. Taken together, our results from metabolomic analyses suggest that there is a modification in the usage of soluble sugars as an energy source to produce ATP.

**Figure 4.**
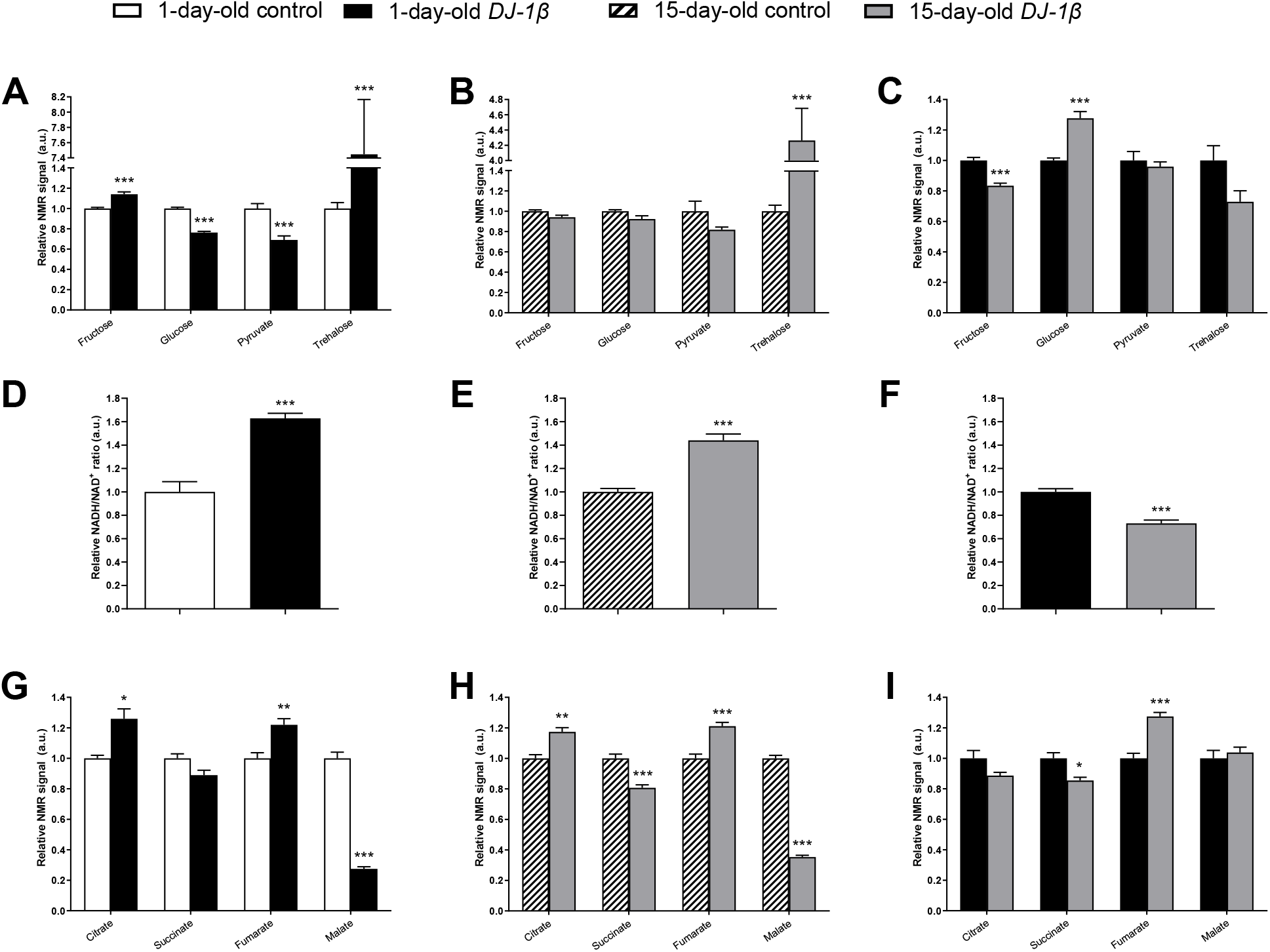
Alterations in carbohydrate content in *DJ-1β* mutant flies. **(A)** Relative NMR signals of selected carbohydrates in 1-day-old *DJ-1β* mutant and control flies. **(B)** Relative NMR signals of selected carbohydrates in 15-day-old *DJ-1β* mutant and control flies. **(C)** Relative NMR signals of selected carbohydrates in 1-day-old and 15-day-old *DJ-1β* mutant flies. **(D)** Relative NADH/NAD^+^ ratio in 1-day-old *DJ-1β* mutants and control flies as an indication of glycolytic pathway activation. **(E)** Relative NADH/NAD^+^ ratio in 15-day-old *DJ-1β* mutants and control flies. **(F)** Relative NADH/NAD^+^ ratio in 1-day-old and 15-day-old *DJ-1β* mutant flies. **(G)** Relative NMR signals of TCA cycle intermediates comparing 1-day-old *DJ-1β* mutants and control flies. **(H)** Relative NMR signals of TCA cycle intermediates comparing 15-day-old *DJ-1β* mutants and control flies. **(I)** Relative NMR signals of TCA cycle intermediates comparing 1-day-old and 15-day-old *DJ-1β* mutant flies.

### Switch from TCA cycle to glycolysis in *DJ-1β* mutant flies

Glycolysis converts glucose to pyruvate, which constitutes an important bridge between this pathway and mitochondrial TCA cycle to produce high amounts of ATP through the electron transport chain (ETC) [46]. In this scenario, NADH/NAD^+^ plays a crucial role in metabolism and highlights the main route to obtain ATP in cells [47]. As shown in Figures 4D-E, both 1-day-old and 15-day-old PD model flies show increased NADH/NAD^+^ ratio when compared to control flies of the same age. This increase is higher in 1-day-old *DJ-1β* mutants (Figure 4F). These results, together with other studies conducted by our group [7], suggest that there is a switch from TCA cycle to glycolysis to obtain ATP. According to this, we observed changes in the NMR signals of TCA cycle intermediates between PD model and control flies (Figures 4G-I). Specifically, 1-day-old and 15-day-old *DJ-1β* mutant flies show an increase in citrate and fumarate and a decrease in malate levels when compared to control flies of the same age (Figures 4G-H). The increase in fumarate levels is higher in 15-day-old *DJ-1β* mutants (Figure 4I). In addition, 15-day-old PD model flies have decreased succinate levels when compared to controls (Figure 4H).

As changes in TCA cycle metabolites were more evident in 15-day-old *DJ-1β* mutants, we decided to focus our studies on this experimental group. We hypothesized that mitochondria become less efficient producing ATP in 15-day-old PD model flies, which was confirmed by measuring ATP levels. As expected, they were reduced in *DJ-1β* mutants when compared to control flies (Figure 5A). This could be explained by an alteration in the TCA cycle. On one hand, citrate is an essential TCA cycle intermediate that acts as a substrate of the Aco enzyme. Aco catalyses the stereospecific isomerization of citrate to isocitrate, and acts as a biosensor of ROS and iron [48]. We observed a decrease in Aco activity in 15-day-old *DJ-1β* mutants compared to control flies (Figure 5B), which may explain the reduction of citrate levels obtained in the metabolomic analysis (Figure 4G-H). On the other hand, increased fumarate, and decreased succinate levels in 15-day-old *DJ-1β* mutants compared to controls could be explained by changes in SDH activity in PD model flies. Mitochondrial SDH oxidizes succinate to fumarate in TCA cycle and reduces ubiquinone in ETC linking both routes [49,50]. Thus, we analysed SDH activity in 15-day-old *DJ-1β* mutants and found that it was decreased when compared to control flies (Figure 5B), therefore suggesting that fumarate must be produced by other ways. In sum, we demonstrated that PD model flies show a shift in metabolic pathway selection to glycolysis due to impairment in TCA cycle.

**Figure 5.**
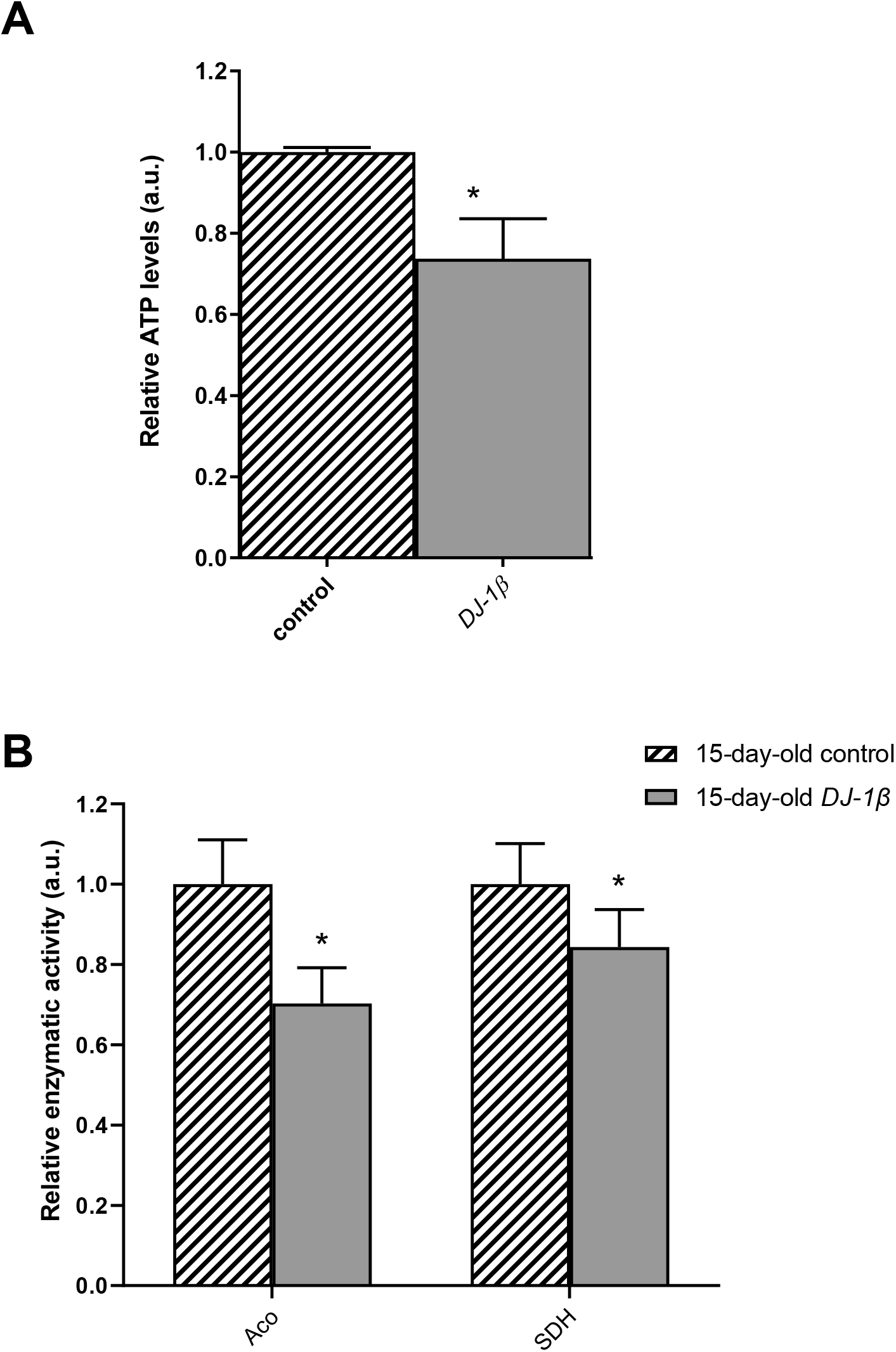
Metabolic switch to glycolysis in 15-day-old *DJ-1β* mutant flies. **(A)** Relative ATP levels in 15-day-old *DJ-1β* mutant flies compared to controls of the same age. **(B)** Relative enzymatic activity of aconitase (Aco) and succinate dehydrogenase (SDH) from the TCA cycle in 15-day-old *DJ-1β* mutant flies compared to controls.

Another possibility to explain the increase in fumarate levels found in 15-day-old *DJ-1β* mutants compared to controls could be an alteration in the UC. Fumarate constitutes a bridge between the TCA cycle and the UC. It is one of the final products of UC, and is transported into the mitochondria where it can be used as a substrate of the TCA cycle (Figure 6A) [51,52]. Our metabolomic analyses showed an increase in fumarate levels but also a reduction of arginine levels in 15-day-old *DJ-1β* mutants. To determine whether these results could be reflecting the existence of changes in the UC in 15-day-old PD model flies, we analysed the expression of the genes encoding the enzymes arginase (*arg*, EC 3.5.3.1) and argininosuccinate lyase (*Argl*, EC 4.3.2.1) (Figure 6A). Arg is a metalloenzyme that catalyses the synthesis of L-ornithine from L-arginine, generating urea [52], while Argl catalyses the reversible conversion of argininosuccinate in L-arginine and fumarate [53]. RT-qPCR analyses revealed an increase in the expression of both genes (*arg* and *Argl*) in *DJ-1β* mutants compared to control flies (Figure 6B). These results are in agreement with changes in fumarate and arginine levels observed in PD model flies and suggest that alterations in the UC could be relevant for PD physiopathology. Further experiments will be required to confirm this assumption.

**Figure 6.**
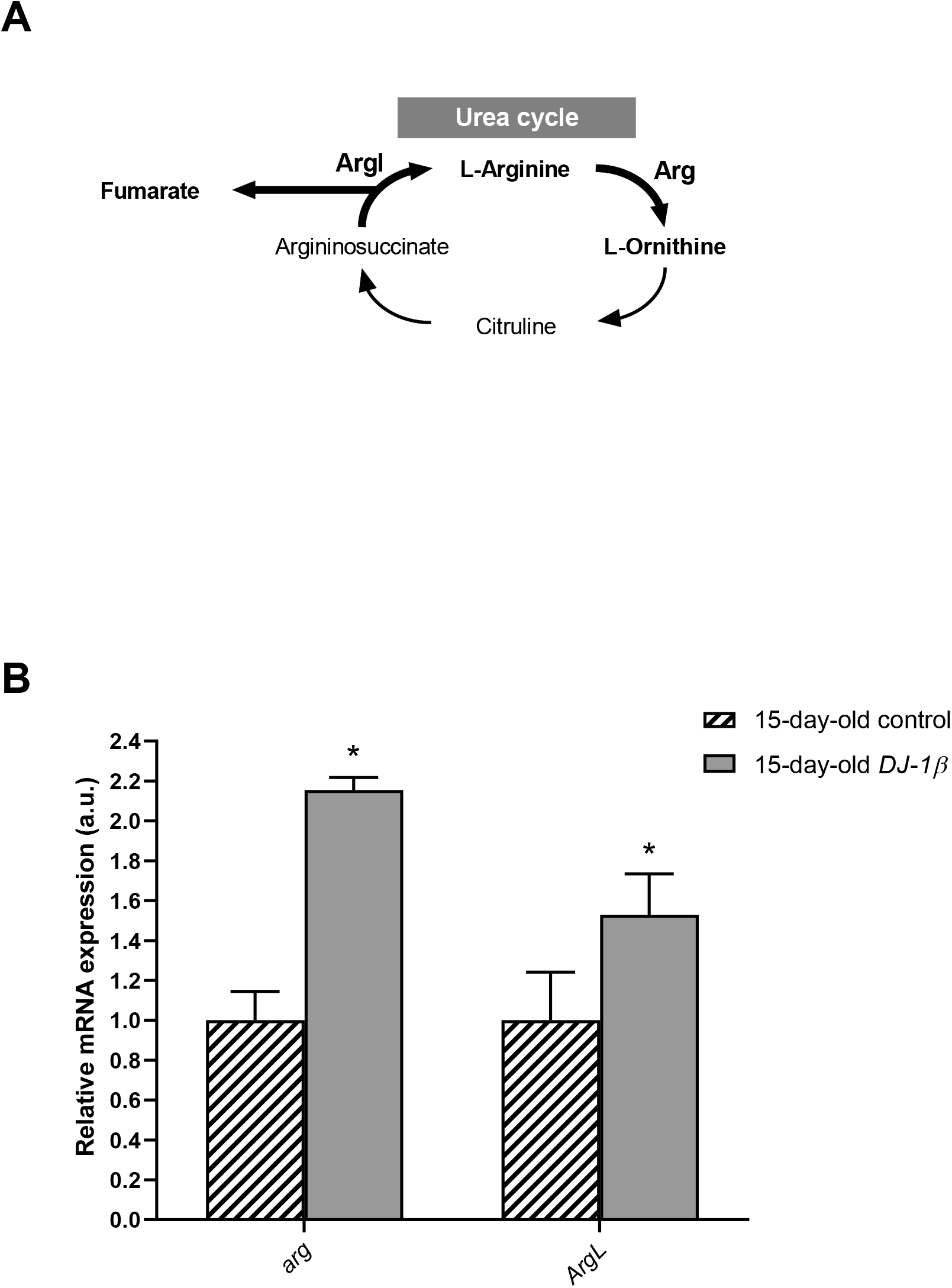
Alterations in the urea cycle in 15-day-old *DJ-1β* mutant flies. (A) Schematic diagram of the urea cycle. Fumarate is a final metabolite that can enter the TCA cycle. (B) Expression levels of selected enzymes of the urea cycle, arginase (*Arg)* and argininosuccinate lyase (*ArgL*).

## DISCUSSION

PD is an incurable and complex disease in which mechanisms causing DA neuronal death are not clear yet [2,5]. Some authors have described a link between redox imbalance, energy failure and metabolic disturbances in PD, leading to the idea that it might be considered a metabolic disease [7,8]. In this study, we used NMR spectroscopy for metabolomic profiling in *DJ-1β* mutant and control flies. The aim of this study was to detect metabolic alterations that might be relevant in PD physiopathology as well as to identify potential therapeutic targets and biomarkers. To date, several metabolomic analyses have been performed in plasm, cerebrospinal fluid or blood of PD patients [43,54–59]. However, human samples are limited to specific tissues and do not allow to get a complete vision of the disease [29–31]. To circumvent this limitation, we have analysed metabolites in the whole body of PD model flies. This experimental approach allows to obtain a complete panorama of PD-associated metabolic defects, which are not restricted to the brain and can influence the physiology of the entire fly [60]. To our knowledge, this is the first metabolomic analysis performed in a *Drosophila* model of fPD. Our results confirm that several metabolic pathways, like amino acid catabolism/anabolism, glycolysis, the TCA cycle, or the UC cycle, might be altered in PD.

Neurons require high levels of energy, which is mainly produced by mitochondria. However, mitochondria become less efficient with age and in several NDs, such as PD, thus leading to a reduction in ATP levels while ROS production increases [46]. In this scenario, amino acids play an important role in the brain, since some of them can be metabolized to fuel cellular energetics when there is insufficient energy [52]. The amino acid profile observed in 1-day-old and 15-day-old *DJ-1β* mutants is consistent with amino acidsanabolism/catabolism imbalance in PD model flies, especially in younger flies (Figure 3). Interestingly, an increase of amino acid catabolism, through transformation to pyruvate and acetyl-CoA or other TCA cycle intermediates to produce energy [61], could compensate ATP deficit due to impaired mitochondrial function. Besides, the reduction in amino acid levels could be also consistent with impairment in protein synthesis in muscles that can exacerbate motor symptoms in PD patients [40].

Among the amino acids whose levels are altered in PD model flies compared to controls, we found BCAAs. BCAAs metabolism/degradation is closely related to carbon metabolism, in particular to glycolysis and the TCA cycle, since they can be transformed to succinyl-CoA that fuels TCA cycle [61]. PD model flies exhibit decreased levels of BCAAs probably due to its degradation to produce energy. A similar situation was found in a metabolomic analysis performed in sebum samples of PD patients [62], thus supporting the validity of the results obtained in our *Drosophila* PD model. Interestingly, a clinical trial (NCT01662414) was performed in PD patients to investigate the effect of Whey protein supplementation, an important source of BCAAs. Results showed that it was able to increase BCAAs and essential amino acid levels, as well as reduced glutathione levels, a key antioxidant that prevents the oxidative damage of DA neurons [41]. In relation to this, a recent study showed that Whey protein supplementation improved motor symptoms in PD patients due to an increase in muscle regeneration [63]. Our results indicate that BCAAs metabolism seems to be decreased in 15-day-old compared to 1-day-old *DJ-1β* mutant flies. A decrease in BCAAs metabolism, especially in aged flies, could enhance PD-related phenotypes as shown in iPD patients [55]. Therefore, BCAAs metabolism could constitute a possible therapeutic target to investigate new PD treatments. Levels of other proteinogenic amino acids are also reduced in *DJ-1β* mutant flies. For example, they present decreased tryptophan levels whose metabolism has been related to neurodegeneration in PD patients at early stages [42,61,64]. Another proteinogenic amino acid affected in *DJ-1β* mutants is glycine, a small amino acid with neuroprotective effects in neuroinflammation, ROS-related damage, and synaptic dysfunction through JNK signalling inactivation [65]. It is likely that a decrease in glycine levels in *DJ-1β* mutants could be contributing to PD pathology. Moreover, there is a significant change in glycine levels when comparing metabolomes of 1-day-old to 15-day-old mutant flies, thus suggesting that this amino acid could be a possible biomarker of PD progression.

PD model flies also exhibit significant increased levels of some non-proteinogenic amino acids compared to controls such as β-alanine. This result could indicate an increase of pyrimidine degradation, as β-alanine is the final product of this pathway. Similar results have been found in *pink1* mutant flies, another *Drosophila* fPD model [66]. Besides, an increase in β-alanine levels causes taurine depletion, which has been related to nerve degeneration [67]. Therefore, alterations in β-alanine metabolism and related pathways could be contributing to PD physiopathology. On the other hand, N-acetyl-aspartate is the most abundant non-proteinogenic amino acid in the central nervous system and a marker of neuronal integrity. It is synthetized in the mitochondria from aspartate and acetyl-CoA, and is then transported to the cytosol, where it is metabolized into aspartate and acetate. Its synthesis is increased in absence of ATP, due to the use of acetate as source of energy [68]. We have found that PD model flies show an important increase in N-acetyl-aspartate (Figure 3) which can be reflecting a deficiency in its metabolism since acetate levels are also decreased in 15-day-old *DJ-1β* mutants compared to control flies of the same age (Table S2).

Alterations in energy metabolism and glucose uptake are associated with physiopathology of several NDs including PD [7,8,52]. Glucose is the main source of energy in the brain and regulation of its metabolism is critical for brain physiology [46]. This is consistent with the observation that *DJ-1β* mutants show dysregulation of carbohydrates metabolism (Figure 4). Supporting this, a decrease of glucose levels in iPSC-derived DA neurons mutant for *PARK2* has been previously observed [69]. In fact, glucose is the sole substrate that can supply the rapid energy demand of neuronal cells through glycolysis [52,70]. Therefore, an increase in glucose consumption through this pathway to restore ATP levels could explain its reduction in our PD model flies [7,71]. Glycolysis is an ATP-producing pathway that could provide considerable amounts of ATP in a less efficient manner than oxidative phosphorylation to support the acute energy demands in neurons [70,72]. DJ-1 has been described to participate in the activity of complex I of ETC binding and stabilizing it [14]. Accordingly, TCA cycle reduction and higher NADH/NAD^+^ ratio in PD model flies suggest the existence of an alteration of this complex and mitochondrial dysfunction, which was also observed in a MPTP-induced PD cell model [73]. Thus, our results show that a shift from TCA cycle to glycolysis is produced in PD model flies [7,37,74]. In addition, it was reported that a reduction of Aco activity causes neurotoxicity in mesencephalic rat cultures, due to the loss of its activity as ROS and iron biosensor [48,75]. Besides, Aco mutant flies exhibited reduced locomotor activity, shortened lifespan, and increased cell death in the developing brain, as well as glycolysis and TCA cycle disturbances that led to decreased ATP levels [48]. Some of the phenotypes are similar to those observed in *DJ-1β* mutant flies [7,27]. On the other hand, reduction of SDH activity could be contributing to lipid accumulation in neurons observed in several NDs, and to produce excitotoxicity, therefore participating in PD pathogenesis and development [76,77]. Thus, our results and these observations suggest that a decrease in the activity of both enzymes might be relevant to PD pathogenesis.

Although TCA cycle activity is decreased in *DJ-1β* mutant flies, there is an increase in some pathway intermediates such as fumarate (Figure 4G-H). TCA cycle metabolites can be produced by other pathways, as could be happening with fumarate and the UC [51]. UC is responsible for the excretion of the nitrogen that cannot be used in amino acid metabolism, and is related to the TCA cycle by fumarate, which is transported into the mitochondria where it can be used as a substrate of Krebs cycle [51,52]. The enhanced expression of *arg* and *Argl* in PD model flies could lead to a general increase of UC activity and to higher fumarate levels, as observed in the metabolomic analysis (Figure 4G-H). Supporting these results, an increase of ArgL activity was observed in a zebrafish PD model based on *DJ-1* deficiency [78]. Changes in *arg* expression levels were also reported to have implications in brain, although its role in this tissue is not known yet [79].. In addition, an enhancement of UC could serve to remove the excess of ammonia caused by the increased amino acid catabolism observed in *DJ-1β* mutant flies, as previously reported in Alzheimer’s disease [79].

In summary, we demonstrate that loss of *DJ-1* function leads to several metabolic alterations, such as amino acid metabolism, carbohydrate metabolism, glycolysis, TCA cycle and UC activity. All of them may be contributing to PD physiopathology and could constitute possible therapeutic targets for this incurable disease. In addition, disturbances in amino acid levels could be potential biomarkers for both early and later stages of PD. Further studies in other preclinical models of fPD and iPD would be required to confirm the results obtained in *DJ-1β* mutant flies.

## Abbreviations

Aco: Aconitase
Arg: Arginase
ArgL: Arginosuccinate lyase
BCAA: branched-chain amino acid
DA: dopaminergic
ETC: electron transport chain
fPD: familial Parkinson’s disease
iPD: idiopathic Parkinson’s disease
ND: neurodegenerative disease
NO: nitric oxide
OPLS-DA: Orthogonal Partial Least Squares Discriminant Analysis
OS: oxidative stress
PCA: Principal Component Analysis
PD: Parkinson’s disease
ROS: reactive oxygen species
SDH: Succinate dehydrogenase
SNpc: *substantia nigra pars compacta*
TCA: tricarboxylic acid
TSP: (Trimethylsilyl)propionic-2,2,3,3-d4 acid sodium
UC: urea cycle

## ACKNOWLEDGMENTS

We are grateful to Dr. Jongkyeong Chung, the Bloomington *Drosophila* Stock Center and the Vienna *Drosophila* Research Center for providing fly stocks.

## Figure legends

**Table S1**. NMR data of selected metabolites from extracts of 1-day-old *DJ-1β* mutant and control flies.

**Table S2**. NMR data of selected metabolites from extracts of 15-day-old *DJ-1β* mutant and control flies.

**Table S3**. NMR data of selected metabolites from extracts of 1-day-old and 15-day-old *DJ-1β* mutant flies

**Table S4**. Results from the pathway enrichment analysis between 1-day-old *DJ-1β* mutants and control flies.

**Table S5**. Selected results from the pathway enrichment analyses in 15-day-old *DJ-1β* mutants and control flies.

**Table S6**. Results from the pathway enrichment analyses in 1-day-oldand 15-day-old *DJ-1β* mutant flies.

