## Supplemental Table 1 for "Metabolic alterations in a *Drosophila* model of Parkinson’s disease based on *DJ-1* deficiency"

**Table S1.** NMR data of selected metabolites from extracts of 1-day-old *DJ-1β* mutant and control flies.

| **Code** | **Metabolite** | **NMR region (ppm)** | | **1-day-old control** | | **1-day-old *DJ-1β*** | | |
| --- | --- | --- | --- | --- | --- | --- | --- | --- |
|  |  | **Right limit** | **Left limit** | **Mean** | **SEM** | **Mean** | **SEM** | **p-value** |
| Var_150 | β-alanine | 3,166 | 3,196 | 1497,021 | 10,324 | 2241,136 | 53,600 | 2,345E-09 |
| Var_45 | Acetate | 1,9 | 1,947 | 1254,178 | 22,332 | 1161,596 | 35,112 | 0,130 |
| Var_254 | Acetyl-aspartate | 2,7 | 2,704 | 4,735 | 0,140 | 11,898 | 0,425 | 1,181E-10 |
| Var_28 | Alanine | 1,469 | 1,505 | 4138,683 | 28,840 | 2749,788 | 85,309 | 2,436E-10 |
| Var_132 | Anserine | 7,078 | 7,105 | 425,873 | 6,873 | 605,140 | 23,402 | 3,273E-05 |
| Var_38 | Arginine | 1,628 | 1,659 | 250,708 | 6,383 | 218,632 | 8,266 | 4,094E-02 |
| Var_70 | Asparagine | 2,922 | 2,928 | 7,376 | 0,276 | 10,465 | 0,689 | 0,008 |
| Var_238 | ATP/ADP | 6,134 | 6,169 | 378,967 | 4,594 | 500,727 | 10,853 | 2,578E-07 |
| Var_261 | Citrate | 2,514 | 2,524 | 28,707 | 0,576 | 36,154 | 1,851 | 0,013 |
| Var_21 | Ethanol | 1,17 | 1,206 | 1296,167 | 99,188 | 1520,882 | 72,816 | 0,210 |
| Var_82 | Formate | 8,455 | 8,468 | 41,417 | 1,579 | 47,178 | 2,238 | 0,151 |
| Var_166 | Fructose | 3,974 | 4,034 | 2339,183 | 24,596 | 2667,412 | 53,791 | 0,001 |
| Var_111 | Fumarate | 6,517 | 6,529 | 18,963 | 0,695 | 23,128 | 0,762 | 0,009 |
| Var_220 | Glucose | 3,709 | 3,716 | 698,250 | 8,449 | 532,351 | 8,781 | 2,402E-09 |
| Var_54 | Glutamine | 2,114 | 2,171 | 1303,499 | 23,227 | 1778,299 | 70,500 | 1,682E-04 |
| Var_151 | Glycerophosphocholine | 3,196 | 3,203 | 204,725 | 5,881 | 119,315 | 3,520 | 1,144E-08 |
| Var_184 | Glycine | 3,557 | 3,57 | 848,584 | 10,408 | 551,874 | 11,357 | 3,371E-12 |
| Var_103 | Guanosine | 8,002 | 8,012 | 20,591 | 0,405 | 14,835 | 0,656 | 2,657E-05 |
| Var_133 | Histidine | 7,005 | 7,019 | 32,760 | 0,646 | 22,541 | 0,872 | 1,081E-06 |
| Var_87 | Hypoxantine | 8,177 | 8,22 | 121,817 | 3,114 | 141,586 | 4,964 | 0,026 |
| Var_7 | Isoleucine | 1,006 | 1,027 | 199,317 | 3,697 | 77,223 | 2,352 | 1,820E-15 |
| Var_5 | Leucine | 0,951 | 0,981 | 674,070 | 16,845 | 264,828 | 5,178 | 7,852E-14 |
| Var_8 | Leucine (triplete) valine (left singulete) | 0,85 | 0,927 | 710,251 | 80,494 | 543,775 | 27,838 | 0,181 |
| Var_195 | Lysine | 3,045 | 3,053 | 156,861 | 6,285 | 50,755 | 4,769 | 2,990E-09 |
| Var_251 | Malate | 2,633 | 2,642 | 47,312 | 1,895 | 12,998 | 0,677 | 3,613E-11 |
| Var_40 | Methionine | 1,703 | 1,769 | 687,468 | 8,824 | 592,802 | 12,880 | 2,990E-04 |
| Var_62 | Methionine-sulfoxide | 2,745 | 2,77 | 1572,816 | 63,483 | 472,262 | 46,353 | 1,448E-09 |
| Var_256 | N-acetyl aspartate | 2,709 | 2,716 | 15,632 | 0,502 | 24,953 | 0,801 | 5,356E-07 |
| Var_149 | NAD+ | 4,478 | 4,502 | 59,812 | 1,621 | 60,424 | 1,586 | 0,850 |
| Var_165 | NADH | 4,203 | 4,239 | 137,540 | 10,262 | 232,759 | 9,099 | 6,565E-05 |
| Var_273 | O-phosphocholine | 3,219 | 3,242 | 5324,258 | 56,353 | 7041,547 | 245,812 | 8,252E-05 |
| Var_117 | Phenylalanine | 7,415 | 7,427 | 36,403 | 0,599 | 22,365 | 0,323 | 8,548E-13 |
| Var_121 | Phenylalanine (sing duplete) | 7,375 | 7,387 | 24,617 | 0,436 | 17,137 | 0,632 | 6,416E-07 |
| Var_162 | Phosphocholine | 4,155 | 4,19 | 664,400 | 11,139 | 905,857 | 48,016 | 0,002 |
| Var_47 | Proline | 1,961 | 1,966 | 9,090 | 0,200 | 8,043 | 0,224 | 0,022 |
| Var_230 | Pyruvate | 2,368 | 2,382 | 944,063 | 45,551 | 651,117 | 38,027 | 0,002 |
| Var_59 | Succinate | 2,398 | 2,417 | 746,371 | 22,023 | 663,889 | 24,254 | 0,089 |
| Var_279 | Threonine | 1,334 | 1,35 | 593,651 | 11,041 | 329,348 | 5,580 | 4,256E-13 |
| Var_309 | Trehalose | 5,185 | 5,214 | 225,223 | 13,010 | 1676,976 | 162,136 | 2,373E-06 |
| Var_99 | Tryptophane | 7,745 | 7,752 | 11,078 | 0,224 | 9,827 | 0,500 | 0,121 |
| Var_9 | Valine | 0,928 | 0,934 | 45,202 | 1,676 | 26,562 | 0,438 | 1,341E-07 |

Note: For each peak we indicate the **code** (variable number for identified metabolites), the integration range (NMR region), the mean of the twelve experimental replicates, and the standard error of the mean (SEM). In all cases we also indicate the statistical significance value (p-value) of the comparison to the corresponding control. All means are highlighted in blue, significant differences (*P*<0.05) are highlighted in red.
