## Supplemental Table 4 for "Metabolic alterations in a *Drosophila* model of Parkinson’s disease based on *DJ-1* deficiency"

**Table S4.** . Results from the pathway enrichment analysis between 1-day-old *DJ-1β* mutants and control flies.

| **Pathway** | **Number of differential metabolites/totals** | **Raw p-value** | **p-value FDR corrected** | **Impact** |
| --- | --- | --- | --- | --- |
| beta-Alanine metabolism | 2/14 | 4,21E-15 | 1,43E-13 | 0,28 |
| Pantothenate and CoA biosynthesis | 2/18 | 4,82E-13 | 8,20E-12 | 0,00 |
| Histidine metabolism | 2/9 | 7,51E-13 | 8,51E-12 | 0,40 |
| Alanine, aspartate and glutamate metabolism | 6/23 | 2,20E-12 | 1,87E-11 | 0,19 |
| Valine, leucine and isoleucine biosynthesis | 4/8 | 6,40E-12 | 4,35E-11 | 0,00 |
| Valine, leucine and isoleucine degradation | 3/38 | 1,99E-11 | 1,13E-10 | 0,00 |
| Aminoacyl-tRNA biosynthesis | 15/48 | 4,03E-11 | 1,96E-10 | 0,00 |
| Starch and sucrose metabolism | 1/14 | 1,21E-10 | 5,14E-10 | 0,01 |
| Glyoxylate and dicarboxylate metabolism | 6/24 | 4,07E-10 | 1,54E-09 | 0,17 |
| Phenylalanine, tyrosine and tryptophan biosynthesis | 1/4 | 1,61E-09 | 4,98E-09 | 0,50 |
| Phenylalanine metabolism | 1/7 | 1,61E-09 | 4,98E-09 | 0,38 |
| Propanoate metabolism | 2/21 | 3,26E-09 | 9,25E-09 | 0,00 |
| Glutathione metabolism | 1/26 | 3,84E-09 | 9,33E-09 | 0,09 |
| Porphyrin and chorophyll metabolism | 1/24 | 3,84E-09 | 9,33E-09 | 0,00 |
| Glycine, serine and threonine metabolism | 3/30 | 1,35E-08 | 2,90E-08 | 0,33 |
| Pyruvate metabolism | 4/22 | 1,36E-08 | 2,90E-08 | 0,28 |
| Glycerophospholipid metabolism | 2/32 | 1,73E-08 | 3,36E-08 | 0,11 |
| Lysine degradation | 1/21 | 1,15E-07 | 2,06E-07 | 0,00 |
| Biotin metabolism | 1/10 | 1,15E-07 | 2,06E-07 | 0,00 |
| Citrate cycle (TCA cycle) | 5/20 | 9,20E-07 | 1,56E-06 | 0,24 |
| Pyrimidine metabolism | 2/40 | 2,93E-06 | 4,74E-06 | 0,00 |
| Purine metabolism | 3/63 | 4,20E-06 | 6,50E-06 | 0,02 |
| Glycolysis/Gluconeogenesis | 4/26 | 3,25E-05 | 4,81E-05 | 0,13 |
| Arginine biosynthesis | 3/12 | 7,25E-05 | 1,03E-04 | 0,40 |
| Cysteine and methionine metabolism | 2/32 | 2,29E-04 | 3,11E-04 | 0,14 |
| D-Glutamine and D-glutamate metabolism | 1/5 | 4,54E-04 | 5,72E-04 | 0,00 |
| Nitrogen metabolism | 1/5 | 4,54E-04 | 5,72E-04 | 0,00 |
| Tyrosine metabolism | 2/33 | 5,15E-04 | 6,26E-04 | 0,04 |
| Taurine and hypotaurine metabolism | 1/7 | 4,72E-03 | 5,48E-03 | 0,20 |
| Arginine and proline metabolism | 2/31 | 4,84E-03 | 5,48E-03 | 0,17 |
| Amino sugar and nucleotide sugar metabolism | 1/34 | 3,12E-02 | 3,42E-02 | 0,00 |
| Tryptophan metabolism | 1/30 | 1,07E-01 | 1,14E-01 | 0,21 |
| Butanoate metabolism | 1/14 | 1,11E-01 | 1,14E-01 | 0,00 |
| Nicotinate and nicotinamide metabolism | 1/9 | 9,52E-01 | 9,52E-01 | 0,37 |
| D-Glutamine and D-glutamate metabolism | 1/5 | 4,54E-04 | 5,72E-04 | 0,00 |

Note: **Number of differential metabolites/total** indicates matched number of metabolites in the total number of compounds in the pathway; **raw p-value** is the original p-value calculated from the enrichment analysis; **p-value FDR corrected** is the p-value adjusted using False Discovery Rate; the **impact** is the pathway impact value calculated from pathway topology analysis. Significant differences (p-value FDR corrected<0.05) are highlighted in blue; high impact scores among significant pathways are highlighted in yellow.
