## Supplemental Table 5 for "Metabolic alterations in a *Drosophila* model of Parkinson’s disease based on *DJ-1* deficiency"

**Table S5.** Selected results from the pathway enrichment analyses in 15-day-old *DJ-1β* mutants and control flies.

| **Pathway** | **Number of differential metabolites/totals** | **Raw p-value** | **p-value FDR corrected** | **Impact** |
| --- | --- | --- | --- | --- |
| Glyoxylate and dicarboxylate metabolism | 6/24 | 1,03E-11 | 3,50E-10 | 0,17 |
| beta-Alanine metabolism | 2/14 | 8,98E-11 | 1,53E-09 | 0,28 |
| Glycerophospholipid metabolism | 2/32 | 4,74E-10 | 4,15E-09 | 0,11 |
| Citrate cycle (TCA cycle) | 5/20 | 4,89E-10 | 4,15E-09 | 0,24 |
| Pantothenate and CoA biosynthesis | 2/18 | 2,63E-09 | 1,79E-08 | 0,00 |
| Taurine and hypotaurine metabolism | 1/7 | 5,08E-09 | 2,88E-08 | 0,20 |
| Histidine metabolism | 2/9 | 2,79E-08 | 1,36E-07 | 0,40 |
| Propanoate metabolism | 2/21 | 1,15E-07 | 4,89E-07 | 0,00 |
| Pyruvate metabolism | 4/22 | 1,88E-07 | 7,09E-07 | 0,28 |
| Lysine degradation | 1/21 | 7,13E-07 | 2,20E-06 | 0,00 |
| Biotin metabolism | 1/10 | 7,13E-07 | 2,20E-06 | 0,00 |
| Valine, leucine and isoleucine biosynthesis | 4/8 | 1,24E-06 | 3,52E-06 | 0,00 |
| Starch and sucrose metabolism | 1/14 | 1,52E-06 | 3,98E-06 | 0,01 |
| Aminoacyl-tRNA biosynthesis | 15/48 | 2,13E-06 | 5,17E-06 | 0,00 |
| Valine, leucine and isoleucine degradation | 3/38 | 3,20E-06 | 7,25E-06 | 0,00 |
| Pyrimidine metabolism | 2/40 | 6,35E-06 | 1,35E-05 | 0,00 |
| Alanine, aspartate and glutamate metabolism | 6/23 | 1,33E-05 | 2,66E-05 | 0,19 |
| Glutathione metabolism | 1/26 | 1,74E-05 | 3,11E-05 | 0,09 |
| Porphyrin and chlorophyll metabolism | 1/24 | 1,74E-05 | 3,11E-05 | 0,00 |
| Glycine, serine and threonine metabolism | 3/30 | 3,65E-05 | 6,20E-05 | 0,33 |
| Arginine biosynthesis | 3/12 | 2,89E-04 | 4,67E-04 | 0,40 |
| Nicotinate and nicotinemide metabolism | 1/9 | 4,03E-04 | 6,23E-04 | 0,37 |
| Tyrosine metabolism | 2/33 | 2,52E-03 | 3,72E-03 | 0,04 |
| D-Glutamine and D-Glutamate metabolism | 1/5 | 7,31E-03 | 9,94E-03 | 0,00 |
| Nitrogen metabolism | 1/5 | 7,31E-03 | 9,94E-03 | 0,00 |
| Purine metabolism | 3/63 | 7,64E-03 | 9,98E-03 | 0,02 |
| Glycolysis/Gluconeogenesis | 4/26 | 1,76E-02 | 2,21E-02 | 0,13 |
| Amino sugar and nucleotide sugar metabolism | 1/34 | 9,32E-02 | 1,13E-01 | 0,00 |
| Cysteine and methionine metabolism | 2/32 | 1,86E-01 | 2,18E-01 | 0,14 |
| Arginine and proline metabolism | 2/31 | 2,56E-01 | 2,90E-01 | 0,17 |
| Phenylalanine, tyrosine and tryptophan biosynthesis | 1/4 | 3,06E-01 | 3,25E-01 | 0,50 |
| Phenylalanine metabolism | 1/7 | 3,06E-01 | 3,25E-01 | 0,38 |
| Butanoate metabolism | 1/14 | 4,49E-01 | 4,63E-01 | 0,00 |
| Tryptophan metabolism | 1/30 | 9,60E-01 | 9,60E-01 | 0,21 |
| Glyoxylate and dicarboxylate metabolism | 6/24 | 1,03E-11 | 3,50E-10 | 0,17 |

Note: In particular, **number of differential metabolites/total** indicates matched number of metabolites in the total number of compounds in the pathway; **raw p-value** is the original p-value calculated from the enrichment analysis; **p-value FDR corrected** is the p-value adjusted using False Discovery Rate; the **impact** is the pathway impact value calculated from pathway topology analysis. Significant differences (p-value FDR corrected<0.05) are highlighted in blue; high impact scores among significant pathways are highlighted in yellow.
