## Supplemental Table 6 for "Metabolic alterations in a *Drosophila* model of Parkinson’s disease based on *DJ-1* deficiency"

**Table S6.** Results from the pathway enrichment analyses in 1-day-old and 15-day-old *DJ-1β* mutant flies.

| **Pathway** | **Number of differential metabolites/totals** | **Raw p-value** | **p-value FDR corrected** | **Impact** |
| --- | --- | --- | --- | --- |
| Glycerophospholipid metabolism | 2/32 | 1,85E-05 | 6,29E-04 | 0,11 |
| Alanine, aspartate and glutamate | 6/23 | 1,98E-04 | 3,36E-03 | 0,19 |
| Valine, leucine and isoleucine degradation | 3/38 | 2,27E-03 | 1,93E-02 | 0,00 |
| Amino sugar and nucleotide sugar metabolism | 1/34 | 2,27E-03 | 1,93E-02 | 0,00 |
| Valine, leucine and isoleucine biosynthesis | 4/8 | 3,28E-03 | 2,23E-02 | 0,00 |
| Citrate cycle (TCA cycle) | 5/20 | 4,54E-03 | 2,57E-02 | 0,24 |
| Aminoacyl-tRNA biosynthesis | 15/48 | 6,99E-03 | 2,87E-02 | 0,00 |
| Phenylalanine, tyrosine and tryptophan biosynthesis | 1/4 | 7,60E-03 | 2,87E-02 | 0,50 |
| Phenylalanine metabolism | 1/7 | 7,60E-03 | 2,87E-02 | 0,38 |
| Histidine metabolism | 2/9 | 1,75E-02 | 5,83E-02 | 0,40 |
| Tyrosine metabolism | 2/33 | 1,89E-02 | 5,83E-02 | 0,04 |
| Butanoate metabolism | 1/14 | 2,23E-02 | 6,32E-02 | 0,00 |
| Arginine biosynthesis | 3/12 | 2,82E-02 | 7,38E-02 | 0,40 |
| Glutathione metabolism | 1/26 | 4,62E-02 | 1,05E-01 | 0,09 |
| Porphyrin and chlorophyll metabolism | 1/24 | 4,62E-02 | 1,05E-01 | 0,00 |
| Propanoate metabolism | 2/21 | 6,00E-02 | 1,27E-01 | 0,00 |
| Pyruvate metabolism | 4/22 | 1,05E-01 | 1,99E-01 | 0,28 |
| Starch and sucrose metabolism | 1/14 | 1,08E-01 | 1,99E-01 | 0,01 |
| Glycolysis/Gluconeogenesis | 4/26 | 1,11E-01 | 1,99E-01 | 0,13 |
| Nicotinate and nicotinamide metabolism | 1/9 | 1,25E-01 | 2,11E-01 | 0,37 |
| Pantothenate and CoA biosynthesis | 2/18 | 1,30E-01 | 2,11E-01 | 0,00 |
| Glyoxylate and dicarboxylate metabolism | 6/24 | 1,43E-01 | 2,20E-01 | 0,17 |
| Glycine, serine and threonine metabolism | 3/30 | 2,20E-01 | 3,26E-01 | 0,33 |
| Arginine and proline metabolism | 2/31 | 2,50E-01 | 3,54E-01 | 0,17 |
| Lysine degradation | 1/21 | 3,36E-01 | 4,40E-01 | 0,00 |
| Biotin metabolism | 1/10 | 3,36E-01 | 4,40E-01 | 0,00 |
| Tryptophan metabolism | 1/30 | 4,21E-01 | 5,20E-01 | 0,21 |
| Taurine and hypotaurine metabolism | 1/7 | 4,30E-01 | 5,20E-01 | 0,20 |
| Pyrimidine metabolism | 2/40 | 4,43E-01 | 5,20E-01 | 0,00 |
| beta-Alanine metabolism | 2/14 | 5,36E-01 | 6,07E-01 | 0,28 |
| D-Glutamine and D-glutamate metabolism | 1/5 | 6,16E-01 | 6,55E-01 | 0,00 |
| Nitrogen metabolism | 1/5 | 6,16E-01 | 6,55E-01 | 0,00 |
| Purine metabolism | 3/63 | 7,08E-01 | 7,30E-01 | 0,02 |
| Cysteine and methionine metabolism | 2/32 | 9,08E-01 | 9,08E-01 | 0,14 |
| Glycerophospholipid metabolism | 2/32 | 1,85E-05 | 6,29E-04 | 0,11 |

Note: In particular, **number of differential metabolites/total** indicates matched number of metabolites in the total number of compounds in the pathway; **raw p-value** is the original p-value calculated from the enrichment analysis; **p-value FDR corrected** is the p-value adjusted using False Discovery Rate; the **impact** is the pathway impact value calculated from pathway topology analysis. Significant differences (p-value FDR corrected<0.05) are highlighted in blue; high impact scores among significant pathways are highlighted in yellow.
